# Language of Stains: Tokenization Enhances Multiplex Immunofluorescence and Histology Image Synthesis

**DOI:** 10.1101/2025.03.04.641512

**Authors:** Zachary Sims, Sandhya Govindarajan, Gordon Mills, Ece Eksi, Young Hwan Chang

## Abstract

Multiplex tissue imaging (MTI) is a powerful tool in cancer research, allowing spatially resolved, single-cell phenotype analysis. However, MTI platforms face challenges such as high costs, tissue loss, lengthy acquisition times, and complex analysis of large, multichannel images with batch effects. To address these challenges, we propose a novel computational method to model the interactions between dozens of panel markers and Hematoxylin & Eosin (H&E) staining, enabling *in-silico* generation of marker stains. This approach reduces the reliance on experimentally measured markers, bridging low-cost H&E data with MTI’s high-content information. Our approach uses a two-stage framework for channel-wise bioimage synthesis: first, vector quantization learns a visual token vocabulary, then a bidirectional transformer infers missing markers through masked language modeling. Comprehensive bench-marking across different MTI platforms and tissue types demonstrates the effectiveness of our method in improving marker prediction while maintaining biological relevance. This advance makes high-dimensional multiplex tissue imaging more accessible and scalable, supporting deeper insights and potential clinical applications in cancer research.

## 1 Introduction

Multiplex tissue imaging (MTI) [1, 11, 16, 17] is an emerging powerful technology in cancer research, enabling the simultaneous measurement of multiple proteins at single-cell resolution while preserving spatial context. Large consortiums such as HuBMAP [7] and HTAN [22] have widely adopted cellular “atlases” to study cell phenotypes and the tumor microenvironment, with the potential to improve patient outcomes and identify novel therapeutic targets. Despite its promise, MTI faces challenges that limit clinical feasibility, including tissue degradation from iterative staining cycles, long acquisition times, and high costs [17, 31]. To address these limitations, we propose a computational model that identifies redundant markers that can be removed from the experimental assay and instead predicted *in-silico*, thereby streamlining the MTI workflow.

Several approaches have been proposed for imputing markers from MTI [24, 25, 28, 31]. Ternes *et al*. [28] introduced a multi-encoder variational autoencoder [27] to reconstruct the full multichannel images from a limited subset of channels. Sims *et al*. [24] improved this with a transfomer-based masked autoencoder (MAE) [13], enhancing predictive accuracy with reduced panels and enabling any-to-any mapping between marker channel sets, thus simplifying training and improving model flexibility. Another approach, 7-UP, leverages cellular morphology and a few antibody stains to generate high-dimensional immunofluorescence images [31]. Collectively, these methods advance computational marker imputation, enhancing the scalability and accessibility of multiplex tissue imaging.

Beyond marker imputation, researchers have explored predicting immunofluorescence (IF) stains from Hematoxylin and Eosin (H&E)[4, 5, 23, 30]. However, existing H&E-to-IF models face challenges as image-to-image translation is typically limited to a few markers. This limitation arises because paired H&E and IF datasets are usually obtained from adjacent tissue sections, separated by approximately 5 microns, due to the tissue degradation issue associated with MTI assays such as CyCIF[17]. Emerging MTI platforms, such as the RareCyte Orion system [16], overcome this limitation by enabling the acquisition of up to 19 IF markers in a single round, preserving tissue integrity for subsequent H&E staining on the same section.

In this paper, we focus on MTI imputation through multimodal integration (i.e., MTI and H&E) to enhance marker prediction accuracy. Our key contributions include:

– Introducing a two-stage framework for channel-wise bioimage synthesis, where a channel-independent visual token vocabulary is learned first, followed by a bidirectional transformer for inter-channel relationships, enhancing biomarker image synthesis.

– Identifying an optimally reduced marker set and improving prediction precision by integrating same-slide MTI data and H&E for colorectal cancer, as well as high-plex MTI for prostate cancer.

## 2 Related Work

### 2.1 Vector Quantization

With the increasing adoption of the transformer architecture [29], a key distinction between its applications in computer vision and natural language processing lies in the nature of input representation. While text is inherently discrete due to tokenization, visual data is typically continuous, as seen in Vision Transformer (ViT) [9]. To bridge this gap, Oord *et al*. [19] proposed the Vector Quantized Variational Autoencoder (VQ-VAE), which discretizes images by mapping latent vectors from the encoder to the nearest entries in a fixed-size codebook. The resulting discrete indices are then used by the decoder for image reconstruction. Building on this approach, Esser *et al*. [10] proposed VQGAN, an improved version of VQ-VAE that incorporates perceptual and adversarial losses to improve reconstruction fidelity.

Beyond facilitating language modeling-based approaches, discretizing visual data is also crucial for computational efficiency as directly modeling images in pixel space is highly inefficient. Instead, decomposing image synthesis tasks into distinct visual and semantic compression stages has been shown to significantly improve performance [32].

### 2.2 Masked Generative Modeling

Most generative transformer models adopt the decoder-only, causal attention masking paradigm popularized by GPT [20], where tokens are decoded sequentially, one by one. However, this approach is neither optimal nor efficient for image generation due to the absence of an inherent sequential structure in images [6, 21, 32]. To address this, masked generative modeling has been introduced to remove the inductive biases associated with text sequences. Chang *et al*. [6] proposed MaskGIT, which replaces causal attention with a bidirectional transformer [8], and predicts all tokens simultaneously. To refine the generation process, multiple inference steps are performed, where only the most confidently predicted tokens are retained, while others are re-masked and re-predicted. This parallelized approach significantly accelerates image synthesis compared to autoregressive methods and has also been shown to outperform diffusion models [32].

Given the growing interest in masked generative modeling, extending this approach to multichannel microscopy imaging presents a compelling opportunity. While transformers have been explored for feature extraction in microscopy image [2, 3, 14], their potential for generative tasks remains largely underexplored [24].

## 3 Methods

Building on masked token prediction for image synthesis [6, 15, 32] and masked image modeling for MTI channel imputation [14, 24], we propose a two-stage pipeline as illustrated in Figure 1. First, multichannel images are transformed into discrete “visual tokens” (middle). Second, missing tokens are imputed to reconstruct absent MTI channels (right).

**Fig. 1.**
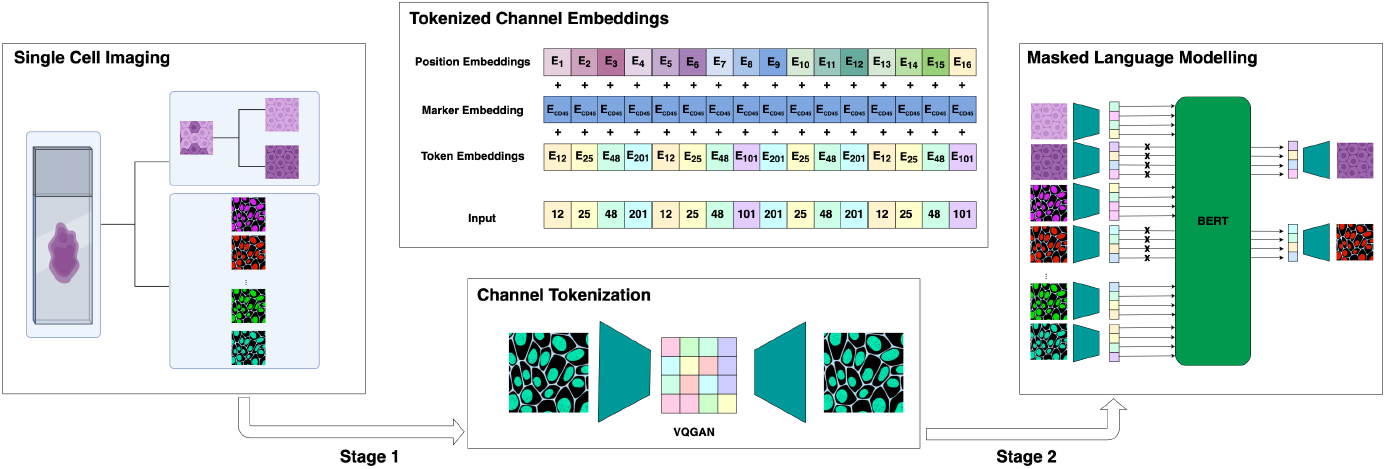
Framework for modeling MTI and H&E as visual language. **Left:** input consists of 64 *×* 64 single-channel cell crops from whole slide images. **Middle:** VQGAN discretizes image channels into 16-integer sequences, treating each channel as a segment like BERT [8]. **Right:** a Masked Language Model predicts discretized masked channels.

### 3.1 Channel Tokenizer

To transform single-cell MTI images into a sequence of discrete tokens, we adopt the MaskGIT approach [6] using the original VQGAN implementation [10] with a codebook size of 256. For channel imputation, our goal is to tokenize an image 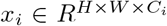 such that we can later input an image with a reduced number of channels, *C*_*k*_, where *C*_*i*_ *> C*_*k*_. To address this discrepancy, we treat *x*_*i*_ as *C*_*i*_ single channel images in *R*^*H×W*^. This approach ensures a consistent tokenizer across varying channel numbers without the need for placeholder channels.

By treating each channel independently, the tokenizer learns morphology- and intensity-based features that are not specific to any single IF marker. This shared codebook strategy enhances the model’s ability to capture meaningful patterns and relationships among IF markers more effectively.

### 3.2 Masked Language Model and Panel Selection

For masked language modeling, we follow the BERT [8] setup, specifically using the RoBERTA configuration from HuggingFace [18]. A single multichannel image is tokenized using our pretrained VQGAN, generating a sequence of length *C* · 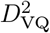, where *C* is the number of input image channels, and *D*_VQ_ represents the dimension of the quantized input image, defined as *D/f*, with *D* as the original input image dimension and *f* as the downscaling factor.

To maintain the analogy with language modeling, we treat the input sequence as a text passage, where each channel functions as a separate sentence. Therefore, we apply the same three embedding layers as in the original BERT implementation [8]: token embeddings, position embeddings, and segment embeddings, as illustrated in Figure 1 (middle).

To determine the optimal set of input markers, we utilize the Iterative Panel Selection (IPS) algorithm from [24]. This greedy selection method prioritizes the “most informative” marker first, specifically the channel *c*_*i*_ that best predicts the remaining channels *C* \*c*_*i*_ when used as a model input. The IPS procedure is formally defined as follows:

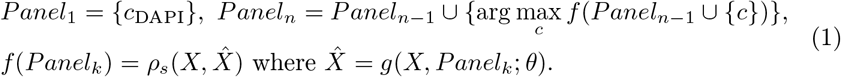

where we set the 1st-marker panel, *Panel*_1_ to DAPI, as it stains the nuclei and provides essential cell location information. Next, we iterate through the remaining *n* −1 markers to identify the marker that, when included with DAPI, yields the highest Spearman correlation (*ρ*_*s*_) between the real and predicted marker intensities using the imputation model *g* with parameters *θ*.

Building on this approach, we introduce an additional heuristic that prioritizes the selection of the marker “hardest to predict”, as defined in Equation (2). Instead of starting with a fixed 1-marker panel and incrementally adding markers to form an *n* −1 panel, we begin with the full panel of *n* markers and progressively remove the easiest-to-predict markers until only the most challenging marker remains. As in the original IPS, we initialize *Panel*_1_ as {*c*_DAPI_}. We refer to this approach as reverse-IPS (rIPS).

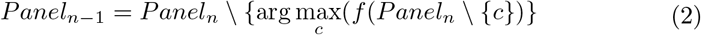

While both heuristics perform similarly, we find that rIPS consistently achieves higher performance when using a low amount of input markers.

## 4 Experimental Data

We evaluate our models on two datasets: a 17-marker Colorectal Cancer (CRC) dataset with same-slide H&E staining from Lin *et al*. [16] and a 40-marker prostate cancer dataset.

For CRC, we split by whole slide image (WSI), using 9 WSIs for training and 1 for testing, covering ∼10.5 million cells with diverse phenotypes. This ensures the model generalizes across tissue regions while avoiding slide-specific biases. The prostate cancer dataset is split by batch (6 for training, 1 for testing) to assess robustness against staining and sample-handling variations, comprising ∼1.6 million cells. Cell segmentation is performed using Mesmer [12], followed by cropping 32 × 32 pixel image patches on each cell to maintain consistent image size and capture the local microenvironment.

For H&E images, we apply color deconvolution to separate the hematoxylin and eosin channels, enhancing the model’s ability to distinguish nuclear and cytoplasmic structures. The resulting two-channel images are treated as additional immunofluorescence (IF) markers in tokenizer training set. To assess potential information loss, we also train a separate tokenizer for H&E using the original RGB format. This follows a multimodal tokenization approach [26], where IF and H&E images map to disjoint token subsets within the model’s token vocabulary (i.e., tokens 0-255 for IF, 256-512 for H&E if our vocabulary size is 512).

## 5 Results

### Model Configurations and Ablations

To optimize model configuration, we evaluate marker imputation across 10 different random marker orders to assess the model’s robustness to marker ordering (Table 1). Our goal is to determine configurations that effectively capture marker expression intensity and structural information while maintaining flexibility in imputing missing markers. Our BERT models use a default setup (12 layers, 12 heads, 768 embedding) and a larger variant (24 layers, 16 heads, 1024 embedding). For MAE, we follow Sims *et al*. [24] with 8 layers encoder/decoder (6 heads, and 1024 embedding) and train a larger MAE of comparable size to the larger BERT model (16 layers, 12 heads, and 1024 embedding).

**Table 1.** Comparison of model configurations. Generative performance is evaluated with 3,6,9, and 12 randomly selected IF markers (H&E adds two channels; “L” denotes a larger model). Top 5 rows show the proposed two-stage model (VQGAN+BERT variants), and bottom 2 rows show MAE. Results are averaged over 10 runs with 10,000 unseen cells; bolded values indicate top performance.

| Model | Spearman Correlation |  |  |  | SSIM |  |  |  |
| --- | --- | --- | --- | --- | --- | --- | --- | --- |
|  | 3 markers | 6 markers | 9 markers | 12 markers | 3 markers | 6 markers | 9 markers | 12 markers |
| IF only | 0.60±0.04 | 0.73±0.03 | 0.82±0.02 | 0.84±0.04 | 0.63±0.01 | 0.69±0.01 | 0.74±0.01 | 0.75±0.01 |
| IF only w/ full ch. mask | 0.69±0.03 | 0.79±0.02 | 0.86±0.02 | 0.87±0.03 | 0.64±0.01 | 0.70±0.01 | 0.74±0.01 | 0.76±0.01 |
| IF only w/ full ch. mask L | 0.70±0.03 | 0.80±0.02 | 0.86±0.02 | 0.87±0.03 | 0.65±0.01 | 0.70±0.01 | 0.74±0.01 | 0.76±0.01 |
| IF+H&E (deconvolved) | 0.72±0.03 | 0.80±0.02 | 0.86±0.02 | 0.87±0.03 | **0.73±0.01** | **0.78±0.01** | **0.81±0.01** | **0.83±0.01** |
| IF+H&E (RGB) | **0.73±0.02** | **0.81±0.02** | **0.87±0.02** | **0.88±0.03** | 0.69±0.01 | 0.74±0.01 | 0.76±0.00 | 0.78±0.01 |
| MAE [24] | 0.64±0.03 | 0.78±0.02 | 0.86±0.02 | 0.87±0.03 | 0.67±0.01 | 0.73±0.01 | 0.77±0.03 | 0.77±0.02 |
| MAE L | 0.66±0.03 | 0.78±0.03 | 0.85±0.02 | 0.87±0.04 | 0.68±0.02 | 0.73±0.01 | 0.78±0.02 | 0.77±0.03 |

To quantify how well the predicted image channels capture marker expression intensity within each cell, we measure the Spearman correlation between real and predicted mean intensity within the center cell region. Models are tested with 3, 6, 9, and 12 random marker subsets. Structural integrity is evaluated using the Structural Similarity Index Measure (SSIM).

Our results show that the proposed two-stage model significantly outperforms the vanilla MAE model from Sims *et al*. [24], particularly when fewer input markers are available. This improvement stems from our full-channel masking strategy, where a random number of channels, *k*, are entirely masked from the full set of *N*_*c*_ channels during training. In this setup, all tokens within each masked channel 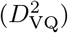 are masked, forcing the model to infer missing information from the unmasked channels. Unlike partial masking, which selects *k* tokens across all 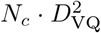 tokens, full-channel masking enhances the model’s ability to capture complex marker relationships, improving imputation performance with limited input markers.

Additionally, incorporating H&E channels as additional inputs further enhances generation quality by providing structural context. These channels complement MTI markers, helping the model learn shared spatial features and produce more realistic imputed channels.

### Optimally Reduced Panel Selection

We use Iterative Panel Selection (IPS) and reverse-IPS (rIPS) with our top-performing two-stage model to identify optimal marker ordering.

Table 2 shows results on the CRC dataset, where IPS and rIPS are applied to 10,000 training cells and evaluated on 10,000 test cells. The optimally reduced panels enhance marker imputation, with our model achieving higher Spearman correlation than MAE model. These heuristics reveal a key trade-off: rIPS yields the best 3-marker panel by selecting distinctive, hard-to-predict markers, making it ideal for minimal yet informative features. IPS, on the other hand, forms the best 12-marker panel by removing redundancy and maximizing predictive performance, making it preferable for broader accuracy. For prostate cancer, our model outperforms vanilla MAE, achieving Spearman correlations of 0.80, 0.86, and 0.93 with 10, 20, and 30 markers as input, improving by 0.01, 0.03, and 0.05, respectively.

**Table 2.** Quantitative comparison of optimally reduced panels for CRC. Generative performance is evaluated using 3,6,9, and 12 IF markers selected via iterative and reverse iterative panel selection on 10,000 training cells. The table presents inference results on 10,000 test cells.

| Model | Panel | Spearman Correlation |  |  |  |
| --- | --- | --- | --- | --- | --- |
|  |  | 3 markers | 6 markers | 9 markers | 12 markers |
| IF+H&E Z=256 w/ full ch. mask | IPS | 0.75 | **0.87** | 0.89 | **0.93** |
| IF+H&E Z=256 w/ full ch. mask | rIPS | **0.77** | 0.84 | 0.88 | 0.90 |
| MAE [24] | IPS | 0.68 | 0.79 | 0.86 | 0.89 |
| MAE | rIPS | 0.69 | 0.80 | 0.86 | 0.91 |

### Marker imputation and relationships

Figure 2 illustrates model-generated MTI images, where 14 channels are predicted from four inputs: DAPI, FOXP3, PD-1, and H&E. Selected via rIPS, these markers improved Spearman correlation by 0.04 over random selection with three inputs. This highlights the model’s ability to infer missing marker patterns while adapting to cell types. The top-left panel shows an Ecad^+^/PanCK^+^ tumor cell, while the top-right features a CD8^+^ T cell (CD3e^+^/CD8a^+^). Figure 2 bottom panel shows an example of a tumor cell in prostate cancer where 34 markers are imputed from six markers as input.

**Fig. 2.**
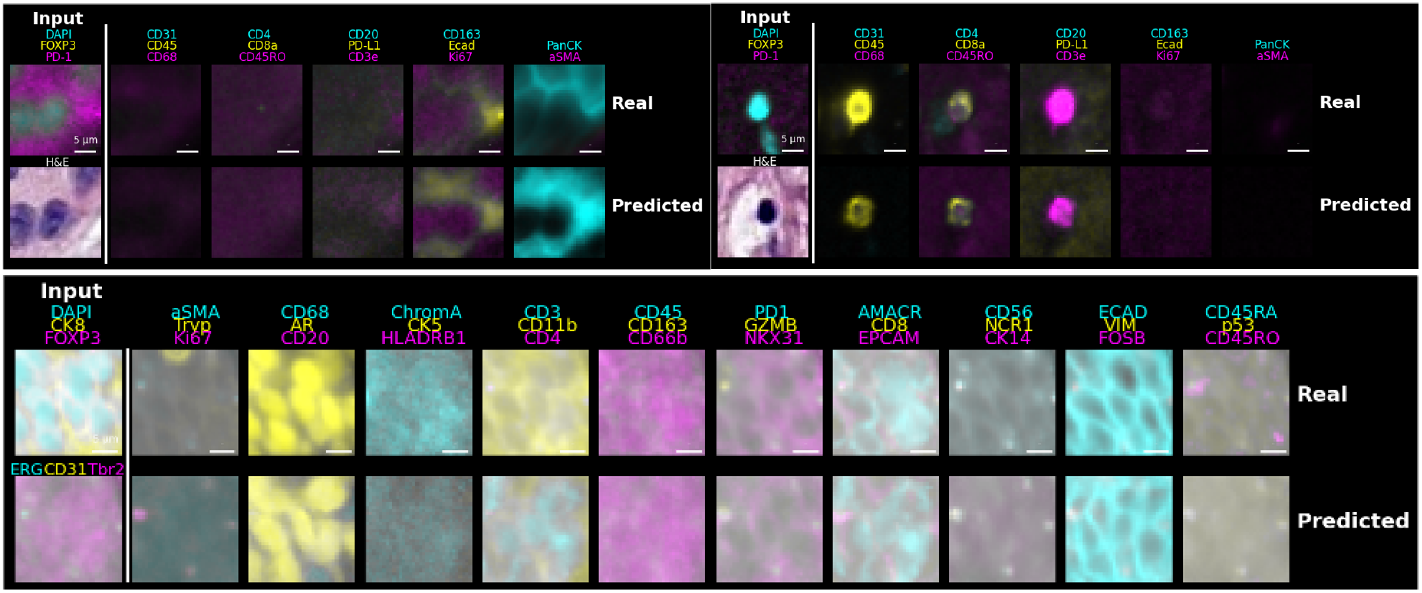
Generation examples using combined H&E and MTI input. The Leftmost column shows input channels (IF markers and/or H&E staining), while the right columns compare ground truth (top row) with predicted image channels (bottom row).

To evaluate how well our model captures biological marker relationships, we compute Cosine similarity between marker embeddings (Figure 3). The left panel shows that while the baseline MAE clusters T cell markers (PD-1, PD-L1, and CD45RO), it fails to associate CD8a with CD3e and CD4, missing key functional relationships. The middle panel demonstrate improvement with VQGAN+BERT, which correctly groups CD3e with CD4 and CD8a, forming a more biologically accurate cluster. In the right panel, adding H&E further refines clustering by linking eosin cytoplasmic markers (*α*SMA and PanCK) and hematoxylin with nuclear markers (Ki67), integrating structural tissue information. Overall, the proposed model, particularly with H&E, offers a more biologically consistent representation, enhancing its ability to infer marker interactions and interpret sub-compartmental, spatial, and morphological features in MTI.

**Fig. 3.**
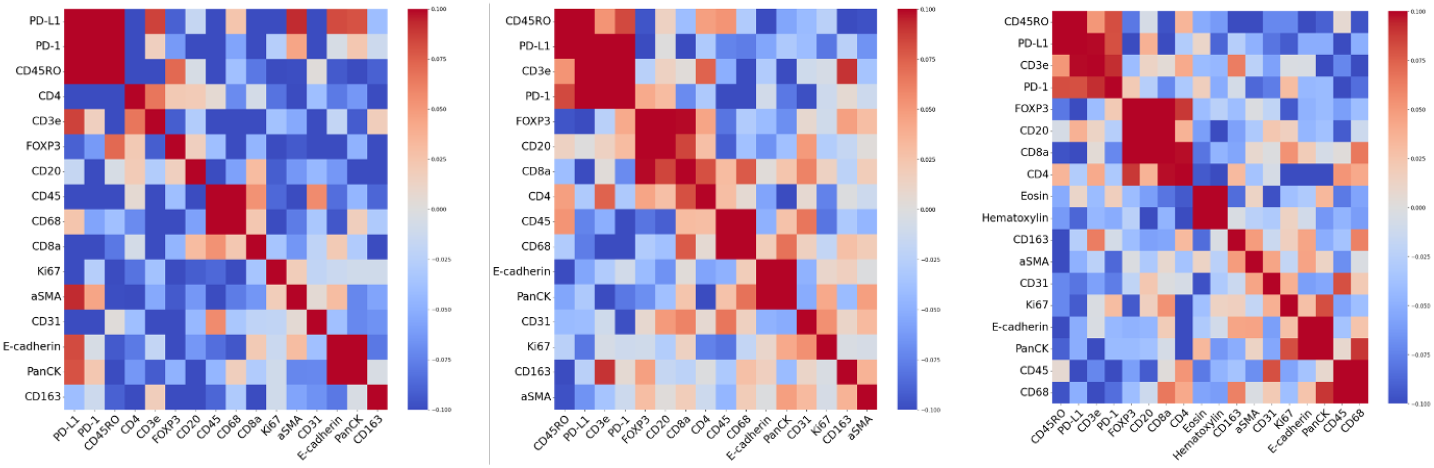
Effect of tokenization on embedding similarities. Left: cosine similarities from the MAE [24] on the CRC dataset. Middle: tokenized language model embedding. Right: Tokenized model with H&E. Pre-tokenizing enhances biologically plausible marker relationships.

## 6 Conclusion

We propose a framework for channel-wise image synthesis of highly multiplex tissue images using vector quantization and bidirectional transformers. This is the first scalable approach to synthesize MTI marker images while integrating MTI and H&E modalities. Our benchmark advances computational biomarker prediction, reducing experimental costs and enabling large-scale spatial tissue analysis.

## Acknowledgments

This work was carried out with major support from the National Cancer Institute (NCI) Human Tumor Atlas Network (HTAN) Research Centers at OHSU (U2CCA233280), and funding (CEDAR3410918) from the Cancer Early Detection Advanced Research Centre at Oregon Health Science University, Knight Cancer Institute (S.E.E.). Y.H.C. is supported by R01 CA253860, R01CA276224, U01 294548, Kuni Foundation Imagination Grants, and the Cancer Early Detection Advanced Research Center (CEDAR) pilot project grant. The resources of the Exacloud high-performance computing environment developed jointly by OHSU and Intel and the technical support of the OHSU Advanced Computing Center are gratefully acknowledged.

